# In the brain of the beholder: bi-stable motion reveals mesoscopic-scale feedback modulation in V1

**DOI:** 10.1101/2024.09.10.612252

**Authors:** Alessandra Pizzuti, Omer Faruk Gulban, Laurentius (Renzo) Huber, Judith Peters, Rainer Goebel

## Abstract

Understanding the neural processes underlying conscious perception remains a central goal in neuroscience. Visual illusions, whether static or dynamic, provide an effective ecological paradigm for studying conscious perception, as they induce subjective experiences from constant visual inputs. While previous neuroimaging studies have dissociated perceptual interpretation of visual motion from sensory input within the motion-sensitive area (hMT+) in humans, less is known about the role of V1 and its relationship to hMT+ during a bistable perception. To address this, we conducted a layer-fMRI study at 7 T with human participants exposed to a bistable motion quartet stimulus. Despite a constant sensory input, the bistable motion quartet elicits switching horizontal and vertical apparent motion percepts likely due to lateral and feedback connections across low and high-level brain regions (feedback processing). As control, we used an “unambiguous” version of the motion quartet, hereafter referred to as “physical” motion stimulus, where horizontal and vertical motion is physically presented as visual stimulus in an alternated fashion (feedforward processing). With the advantage of a sub-millimeter resolution gained at ultra-high field (7 Tesla), we aimed to unveil the differential laminar modulation of V1 (low visual area) and hMT+ (high-visual area) during the physical and bistable condition. Our results indicate that:

1. hMT+ functional activity correlates with conscious perception during both physical and ambiguous stimuli with similar strength. There is no evidence of differential laminar profiles in hMT+ between the two experimental conditions.
2. Between inducer squares, V1 shows a significantly reduced functional response to the ambiguous stimulus compared to the physical stimulus, as it primarily reflects feedback signals with diminished feedforward input. Distinct V1 laminar profiles differentiate the two experimental conditions.
3. The temporal dynamics of V1 and hMT+ become more similar during the ambiguous condition.
4. V1 exhibits reduced specificity to horizontal and vertical motion perception during the ambiguous condition at the retinotopic locations corresponding to the perceived motion.

Our findings demonstrate that during the ambiguous condition, there is a stronger temporal coupling between hMT+ and V1 due to feedback signals from hMT+ to V1. Such feedback to V1 might be contributing to the stabilization of the vivid perception of directed motion at the face of constant ambiguous stimulation.

## 1 Introduction

The ability to process motion information is crucial for human survival, social interaction, spatial navigation and cognitive processing. Understanding how the brain integrates and interprets sensory input to construct a subjective visual experience of moving objects might shed light on fundamental aspects of human perception and cognition. Here, we focus on the ambiguous motion stimuli introduced by Schneider et al. (2019) and investigate how the brain engages in complex neural dynamics within and between early (V1) and high-level visual areas (hMT+) to create an unambiguous conscious perception. The crucial role of high-level visual areas, especially the middle temporal area during motion perception tasks, was repetitively demonstrated in both MT in monkeys and hMT+ in humans. Starting with pioneering electrophysiology experiments in awake, behaving monkeys, higher visual areas were found to be responsible to resolve binocular rivalry tasks, where two incompatible monocular images compete for perceptual dominance (Leopold and Logothetis, 1999; Logothetis, 1998; Logothetis and Schall, 1989); the number of firing neurons indicating the stimulus of the chosen eye was shown to increase progressively across the cortical visual hierarchy: approximately 20% in early areas (V1/V2), around 40% in intermediate areas (MT and V4), and about 90% in later areas (IT) (Polonsky et al., 2000). A similar outcome was demonstrated to occur during the presentation of bistable figures, where selectively neurons in MT (Dodd et al., 2001), medial superior temporal (MST), and parietal cortex (Williams et al., 2003) all show activity reflecting the consciously perceived stimulus. The central role of MT in constructing the perception of motion direction was demonstrated also by the seminal electrical microstimulation study in the monkey during a motion discrimination task (Salzman et al., 1990, 1992). With the advent of fMRI, analogous perceptual phenomena have been studied in living humans, leading to similar results (Muckli et al., 2002; Sterzer et al., 2002). More recently, a high-resolution hMT+ fMRI study at 7T by Schneider et al. (2019) used a bistable motion quartet stimulus that elicits an alternation of horizontal and vertical motion perception during passive viewing, and demonstrated that the experienced percept is associated with increased activity of respective axis-of-motion columnar clusters coding horizontal and vertical motion axes. Here, we use the same experimental paradigm as Schneider et al. (2019) and extended the focus to both hMT+ and V1: based on previous results, we hypothesized that hMT+ resolves the ambiguity of the stimulus and that the feedback loop with V1 serves to stabilize perception. Despite previous attempts (Liu et al., 2019; Lumer and Rees, 1999; Muckli et al., 2005; Supèr et al., 2001; Tong et al., 1998) to understand the neural correlates of conscious perception along the visual hierarchy, the role of V1 and its modulation through feedback signals is still debated in the community. Crucially, feedforward and feedback processing involves different laminar pathways: feedforward projections predominantly terminate in layer IV in sensory cortical areas, whereas feedback projections might terminate in superficial or deep layers (Callaway, 1998, 2004; Felleman and Van Essen, 1991). Arguably high-field fMRI (7 T or higher) with submillimeter voxel resolution (*<*1 iso mm) is the most promising technique to non-invasively differentiate signals from different cortical layers or cortical columns in living humans and potentially disentangle feedback and feedforward processing (Bergmann et al., 2024; De Martino et al., 2018; L. Huber et al., 2017; Kok et al., 2016; Schneider et al., 2019; Shen et al., 2020; Ugurbil, 2016). To understand how information is exchanged between early and high-order visual areas and to test whether a feedback signal from hMT+ targets a specific cortical layer in V1, we recorded 0.8 mm iso-voxel fMRI data at 7 T from nine participants while they perceived the bistable motion quartet stimulation (ambiguous motion condition) (Schneider et al., 2019) (**Figure 1**). While a constant sensory input consisting of two static frames with two inducer squares is presented to the retina (**Figure 1A**), participants perceive directed motion alternating between episodes of right-left (horizontal) and up-down (vertical) motion. The ambiguous motion condition serves as a proxy of a feedback state for V1 (compared to the ‘feedforward’ physical motion condition), especially at non-stimulated locations between inducers. Uniquely, we concurrently defined both columnar clusters in hMT+ and retinotopic clusters reflecting the apparent motion path in V1 and differentiated fMRI signals from different cortical layers. In both areas, we quantified the signal changes occurring during the alternating conscious perception of motion. We are particularly interested in whether feedback from hMT+ modulates upper / lower layers only in retinotopic clusters matching the percept (hypothesis 1) or whether feedback modulates (differentially) both retinotopic clusters (hypothesis 2, see **Figure 1D**). If a participant perceives horizontal / vertical motion, feedback modulation would be observed only in horizontal / vertical retinotopic clusters in V1 according to hypothesis 1, while both retinotopic clusters would be (differentially) modulated for horizontal and vertical percepts according to hypothesis 2. Finally, we discuss whether any source of feedback from hMT+ to the not-matching retinotopic cluster of V1 is the same as the matching retinotopic cluster of V1, i.e., both columnar clusters of hMT+ could potentially contribute to the feedback to the vertical retinotopic cluster of V1 (see two arrows going to vertical (blue) V1 cluster in **Figure 1D**.

**Figure 1:**
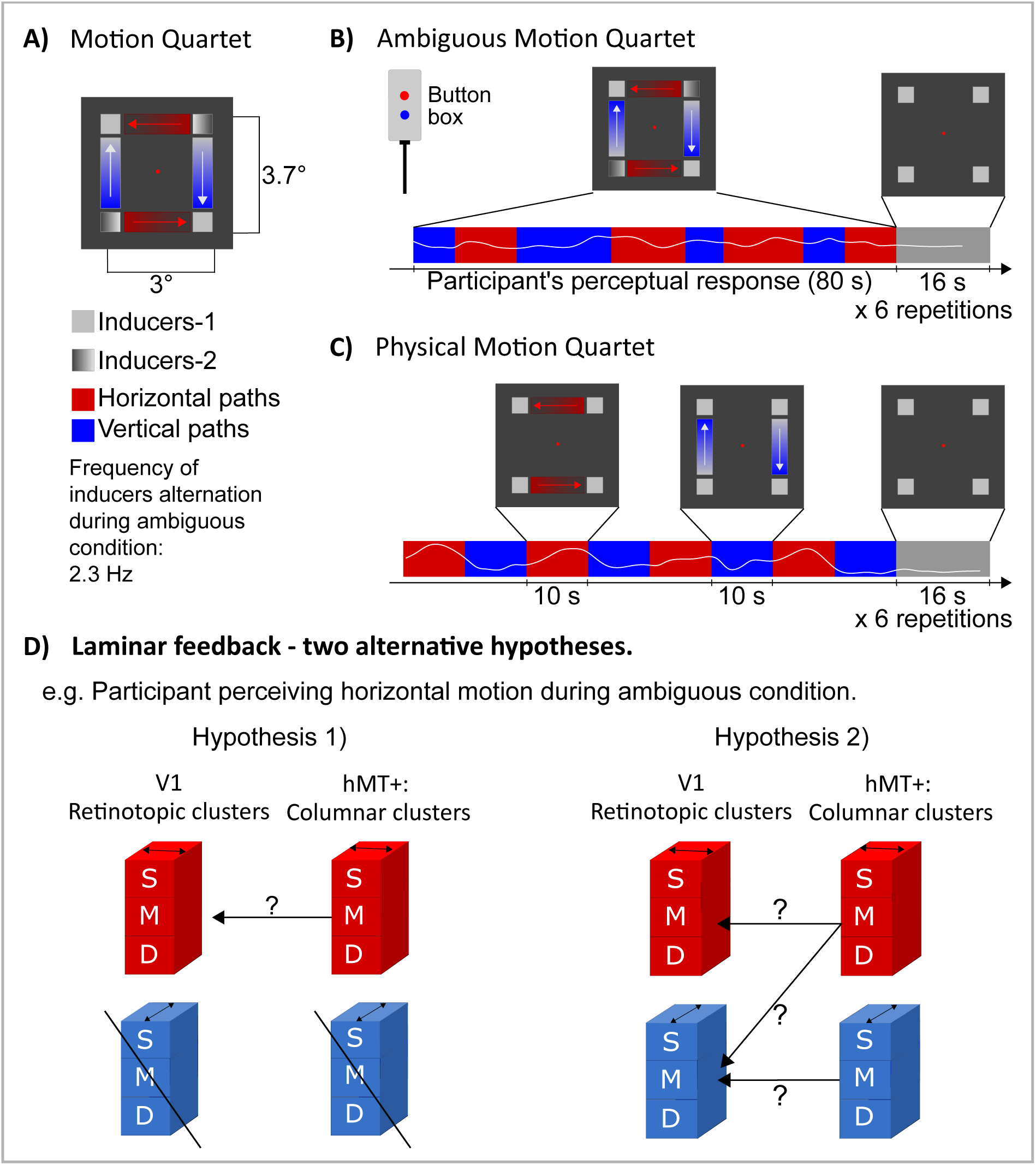
Experiment design. A) Motion quartet stimulus (Animations: https://doi.org/10.6084/m9.figshare. 21908394.v1). B-C) Two versions of the motion quartet stimulus are used in the experiment. (B) During the ambiguous motion the bistable quartet is continuously presented for 80 s to the participant perceiving an alternation of horizontal and vertical motion (participant indicates the percepts by using an MRI-compatible button box during the fMRI session). Colored paths in between inducers are for illustrative purposes only (i.e., not shown during the experiment). (C) During the physical motion quartet participant watches inducers (squares) moving unambiguously horizontally and vertically (10 s) in an alternated fashion. (D) Ambiguous motion condition serves as a proxy for investigating the feedback signal in V1. At mesoscale, we differentiate retinotopic clusters in V1 and motion-specific columnar clusters in hMT+ (horizontal clusters are shown as red blocks, while vertical clusters are shown as blue blocks). Clusters schematically represent a group of neurons that preferentially respond to the horizontal or vertical physical motion condition. Three cortical depths (s: superficial, m:middle, d:deep) coarsely describe the cortical laminar organization. We compare two models of feedback from

## 2 Results

### 2.1 Mesoscopic clusters in hMT+ respond equally strong to physical and ambiguous motion conditions

During the ambiguous motion condition, the participant perceives horizontal and vertical motion in an alternated fashion, similar to the physical motion condition. We aimed to determine whether clusters of voxels that primarily responded to the physical motion condition would also respond during the ambiguous condition. To investigate this, we compared the percent signal changes between the two experimental conditions for each retinotopic/motion cluster in V1 and hMT+ regions separately. Our findings reveal different modulations in these two brain regions. During the ambiguous motion condition, we observed a significant reduction in percent signal change in both retinotopic clusters in V1 (**Figure 2C**). In contrast, the motion clusters in hMT+ exhibited consistent modulation in both conditions (**Figure 2D**). These results align with the expectation that higher-order visual areas, like hMT+, are more engaged than early visual areas, like V1, in resolving ambiguity and creating motion perception (Goebel et al., 1998, 2001; Leopold and Logothetis, 1999; Muckli et al., 2002; Schneider et al., 2019; Sterzer et al., 2002). The reduced modulation in V1 might indicate a response to feedback signals from higher-order areas. Such feedback-driven activity is expected to be lower compared to its feedforward counterpart (Muckli et al., 2005; Shen et al., 2020; Sterzer et al., 2006).

**Figure 2:**
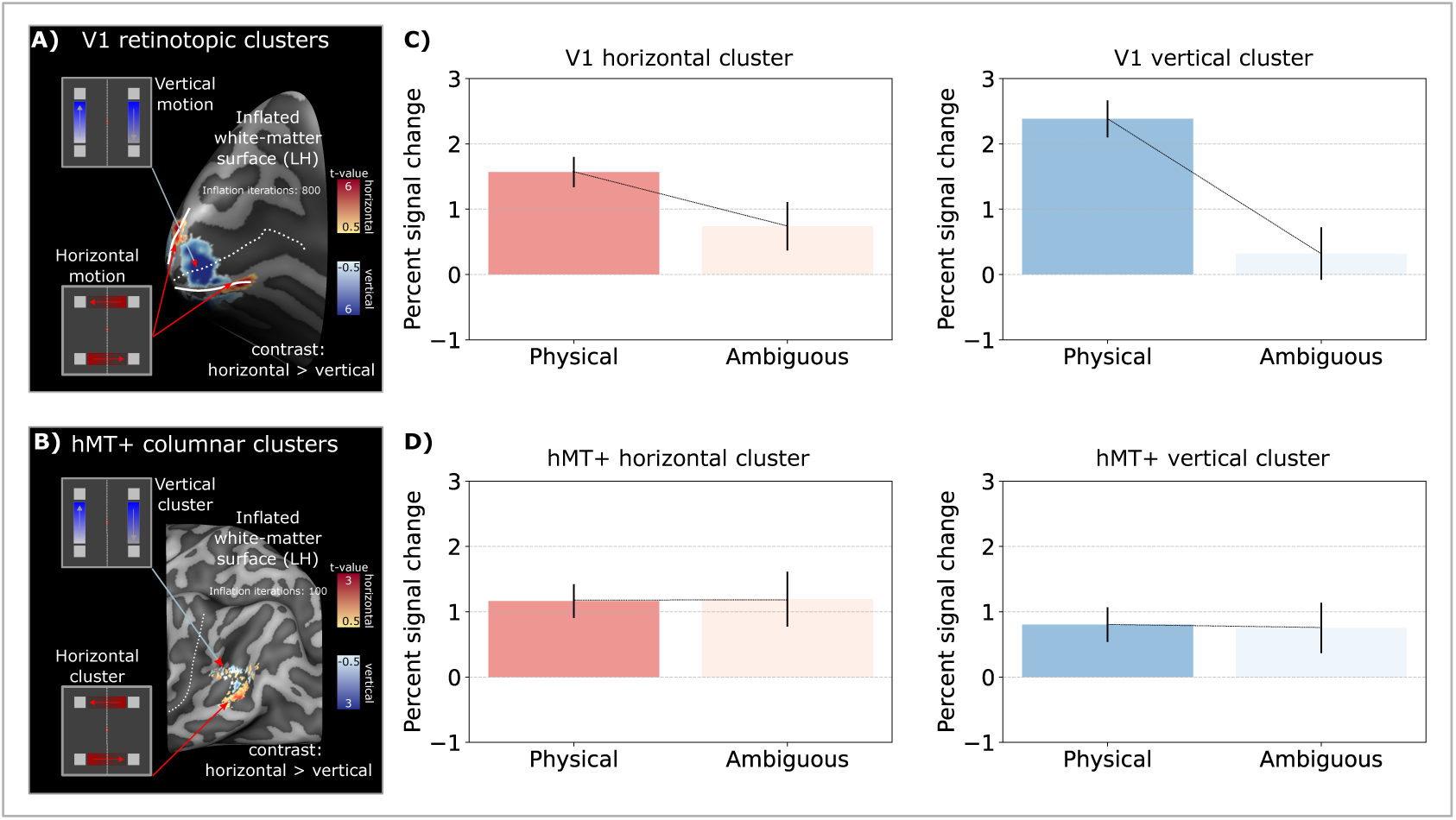
Comparison of V1 and hMT+ fMRI responses during physical and ambiguous motion conditions. (A) V1 retinotopic clusters representing horizontal (in red) and vertical (in blue) motion paths, observed during the physical motion condition, are visualized on the inflated white matter surface. The dotted line delineates the calcarine sulcus, while solid lines indicate V1/V2 boundaries (see **Supplementary Figure 2, 3**). (B) hMT+ horizontal (in red) and vertical (in blue) columnar clusters, defined based on the physical motion condition, are visualized on the inflated white matter surface, with the dotted line indicating the medial temporal sulcus. (C-D) Bar plots display the mean percent signal change (+/-standard error of the mean) computed across participants for both V1 (C) and hMT+ (D) regions, within each cluster (horizontal in red and vertical in blue), for both physical and ambiguous conditions. While hMT+ responses appear invariant between the two experimental conditions, V1 responds substantially weaker during the ambiguous motion condition.

### 2.2 Feedback in V1 during ambiguous motion condition at laminar scale

We investigated whether physical and ambiguous motion conditions were associated with differential feedforward- and feedback-weighted laminar patterns. To this end, we computed laminar profiles for both V1 and hMT+ within each mesoscopic cluster in response to both conditions (**Figure 3**). Our results show that we could successfully distinguish between feedforward and feedback conditions in V1 but not in hMT+. As shown in **Figure 3C**, V1’s laminar profile varies between the two conditions consistently for both horizontal and vertical clusters: while the physical motion condition is characterized by a linear increase towards superficial layers, the ambiguous condition elicits a similar response in all layers. Our findings suggest that the feedback signal received by V1 does not lead to a conventional increase towards superficial layers, that is expected for a feedforward modulation. Since all layers seem to equally contribute to the ambiguous condition, we can’t straightforwardly conclude which layer is receiving the feedback signal. Comparing laminar profiles between two experimental conditions by computing differential laminar profiles is a common approach used to mitigate the unwanted macrovascular signal contribution responsible for the draining vein effect (Aitken et al., 2020; Fracasso et al., 2018; Heynckes et al., 2023; van Mourik et al., 2023 (See our Discussion section for further elaborations)

**Figure 3:**
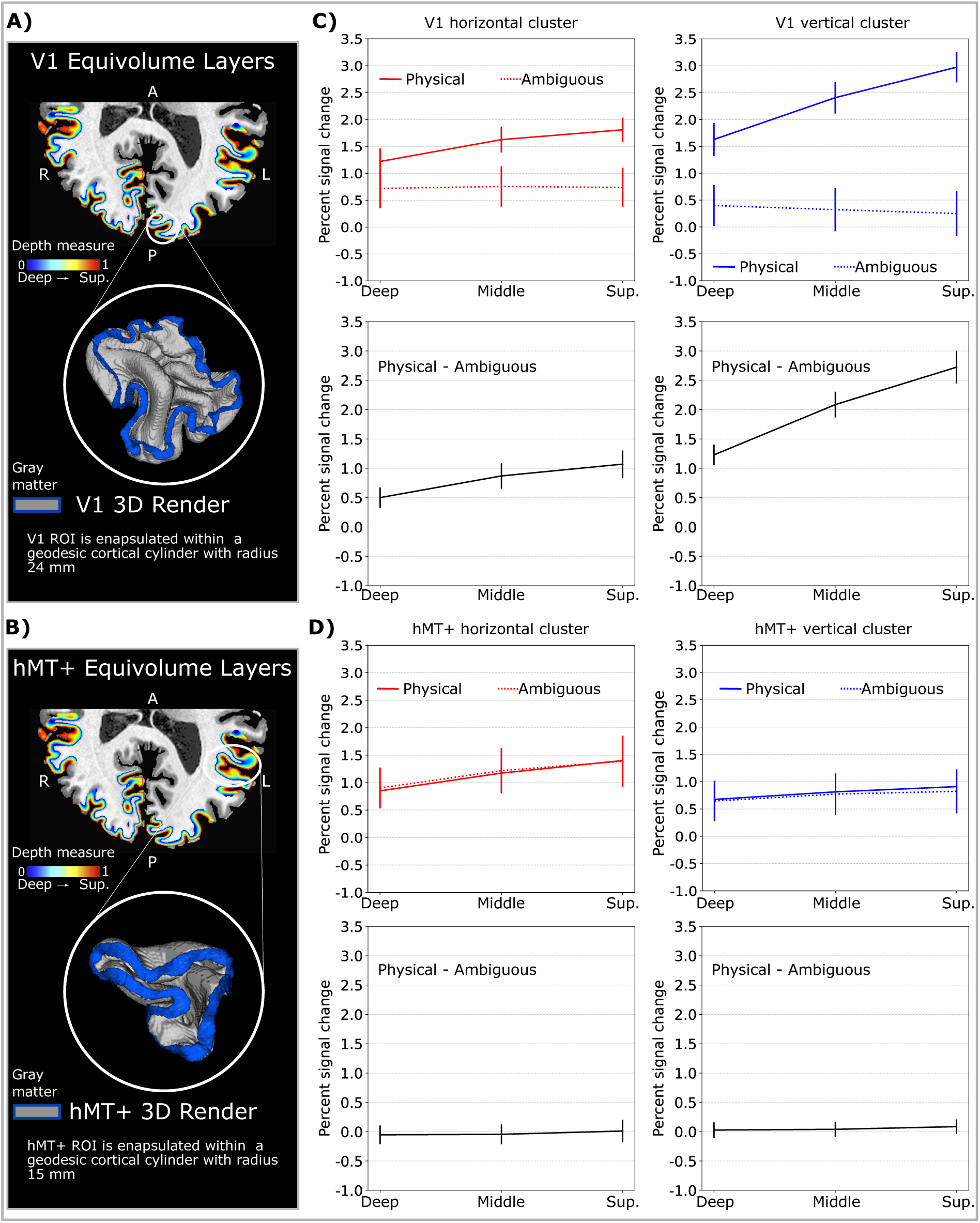
Comparing V1 and hMT+ fMRI response as function of cortical depth as difference between physical and ambiguous motion conditions. A-B) Normalized equivolume-depth measures (in rainbow colors) are computed for each region of interest to characterize the cortical depth and extract layers. White circular shape indicates 3D volumetric rendering of V1 and hMT+ gray matter. C-D) Top row. Group-level laminar profiles for physical motion condition (solid line) and ambiguous motion condition (dotted line) expressed as percent signal change. For each cluster, mean percent-signal change (+/- standard error of the mean) is reported at each of the three cortical layers: deep, middle and superficial. Bottom row. Differential laminar profiles. For each cortical depth, we report the difference between physical and ambiguous

Given our single condition laminar profiles, the differential laminar profiles in V1 resulting from the difference between the percent signal change elicited by the physical and ambiguous conditions (**Figure 3C** **-bottom row**), shows that the strongest difference is consistently observed in the superficial layers for both clusters. This result is a direct consequence of a reduced draining vein effect, as manifested as linear increase, during the ambiguous condition; if we assume that during the feedforward condition V1 laminar profiles are characterized by an expected middle layer modulation (receiving input from the thalamus) plus a linear increase towards pial surface (**Figure 3C** **- top row, solid line**), we can explain V1 laminar profiles during the ambiguous condition as consequence of a redistribution of activity in all layers in response to an absence of middle layer modulation. Laminar BOLD results on feedback conditions without the expected draining vein effect have been already reported (Bergmann et al., 2024; Kok et al., 2016).

Contrary, we didn’t find any laminar differences in hMT+ between the two conditions (**Figure 3D**). The fact that the difference in laminar profiles in hMT+ results in almost zero across all layers (**Figure 3D**, **bottom row**) suggests two possibilities: either the processing in hMT+ does not vary between the two experimental conditions, or the variation exists but is not captured by the current laminar analysis.

### 2.3 Similar temporal dynamics between V1 and hMT+ during ambiguous motion condition

V1 and hMT+ voxels were assigned to horizontal or vertical clusters according to the condition (horizontal or vertical) that elicited the strongest functional response (as t-value) during the physical motion condition. However, the analysis above does not retain the information related to the “strength of the preference”, meaning how much a voxel is preferentially responding to one condition versus the other one. For this reason, we computed a measure of functional specificity for each voxel (as previously established in Pizzuti et al., 2023) of both V1 and hMT+ for each experimental condition. Voxel-wise specificity values were averaged across voxels belonging to the same cluster (**Figure 4A-B**). We found a substantial decrease in specificity only for V1 but not hMT+ clusters from the physical to the ambiguous condition. We investigate this further by reporting the temporal response patterns visualized as time-locked event-related averages (ERA) elicited by V1 and hMT+ clusters during both physical and ambiguous motion conditions (**Figure 4C-D**) while perceiving the preferred (e.g. horizontal cluster response to horizontal motion condition) or not preferred condition (e.g. horizontal cluster response to vertical motion condition). As expected by our voxel’s selection strategy, during the physical motion both V1 and hMT+ are positively activated after the stimulus onset (at 0 s) during the preferred condition. While V1 clusters show a negative hemodynamic response during the not preferred condition, hMT+ clusters show an initial positive peak for 2 s followed by a larger decrease in the not preferred as compared to the preferred condition. During the ambiguous condition, because of the decrease in specificity, we found a positive response from V1 clusters to both preferred and not preferred conditions, with similar temporal dynamics. In contrast, hMT+ event-related averages show an initial positive signal increase for both preferred and not preferred conditions followed by a differential response due to a stronger decrease during the not preferred condition similar as in the physical motion condition. A similar response profile was reported by Schneider et al. (2019). Note that both V1 and hMT+ event-related averages peak around 4 s, suggesting a more synchronous behavior between the two areas that might indicate a common feedback signal or specific feedback from hMT+ to V1.

**Figure 4:**
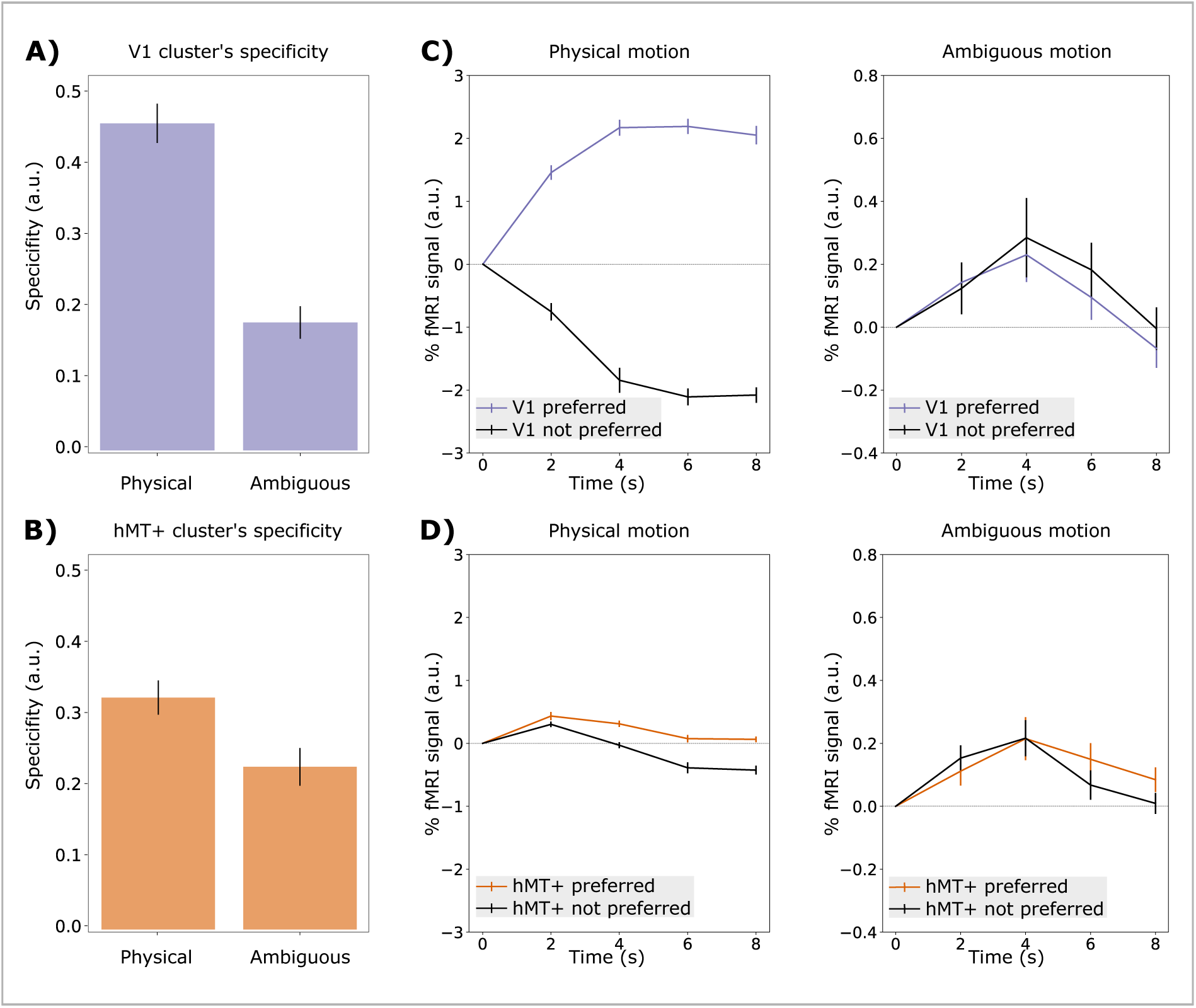
Specificity of V1 and hMT+ clusters (A-B) and event related-averages (C-D) for both physical and ambiguous conditions are shown in separate panels for both V1 and hMT+. (C) V1’s cluster response to not preferred condition only occurs during the ambiguous stimulation and with similar amplitude to the correspondent response to the preferred condition. This double response might indicate that a feedback signal arrives to the clusters in both conditions. (D) In both motion conditions, hMT+’s cluster response (positive initial peak) to not preferred conditions occurs and shows a stronger decrease than the preferred condition.

## 3 Discussion

We used the bistable motion quartet stimulus to investigate motion perception processing in hMT+ and V1 using submillimeter fMRI at 7 T. This allowed us to differentiate functional clusters within each cortical area at laminar resolution. We confirmed that hMT+ clusters maintain functional modulation during the ambiguous motion, similar to the physical condition, with consistent specificity (**Figure 2D**, **Figure 4B-D**). For the first time, we showed that hMT+ cluster organization at the laminar scale remains invariant between conditions (**Figure 3D**). In V1, both horizontal and vertical retinotopic clusters showed a substantially weaker response during ambiguous motion as compared to physical motion condition, with distinct laminar patterns and the strongest changes in superficial layers (**Figure 2C**, **Figure 3C**). We uniquely quantified the cluster’s functional specificity in V1 and hMT+ and demonstrated that similar event-related averages for both preferred and not preferred conditions characterize V1 clusters’ response during the ambiguous motion condition (**Figure 4**). Initially, the hMT+ clusters responded with equal strength followed by a more pronounced decrease in the not preferred condition. The temporal profiles of V1 and hMT+ in the ambiguous condition are strikingly similar with a peak at four seconds suggesting a feedback mechanism between the two areas.

### 3.1 Feedback in V1: origin and traces at mesoscale

In this study, we disentangle feedforward- and feedback-driven processing through experimental manipulation: a “physical” and an “ambiguous” motion condition were used to investigate the role of hMT+ and V1 using layer-fMRI at 7 T. Our results demonstrate that only V1, but not hMT+, shows a new characteristic functional response during the feedback condition compared to the feedforward condition. This fMRI response is characterized by 1) a decreased amplitude (**Figure 2**) and 2) a uniform contribution from different cortical layers (**Figure 3**). This result is in line with the expectation that only a smaller subset of neurons is targeted by the feedback signal, compared to the feedforward condition (Muckli et al., 2005; Shen et al., 2020; Sterzer et al., 2006). Consistently with our results, similar differences between feedforward and feedback conditions have been previously observed by other BOLD laminar-fMRI studies (Aitken et al., 2020; Carricarte et al., 2024; Kok et al., 2016; Lawrence et al., 2018; Thomas et al., 2023). Our results demonstrate that the laminar response in V1 undergoes a different modulation during the feedback condition compared to the physical condition, leading to a redistribution of activity across all layers (**Figure 3C-top row, dotted line**). Even if the strongest (de-)modulation is observed in superficial layers, we can’t conclude that superficial layers are the target of the feedback signals.

Uniquely, we analyzed the temporal dynamic of V1 and hMT+, as event-related averages, during both the physical and the ambiguous motion condition (**Figure 4C-D**) and show a striking temporal similarity between the two areas only during the ambiguous motion condition. This temporal feature might support the hypothesis that a feedback signal from hMT+ to V1, only occurring during the bistable condition, comes with an increase in synchrony between the two communicating areas.

Furthermore, we demonstrated that during a perceptual state (e.g. horizontal motion), beside the V1 voxels coding for the correspondent retinotopic location (horizontal cluster) also the voxels coding for the alternative retinotopic location (vertical cluster) elicit a positive signal (**Figure 4**). We speculate that this is due to a global feedback mechanism from hMT+ to V1, and that the feedback to the not-matching retinotopic cluster in V1 (e.g., V1 retinotopic vertical cluster) might be attributed to two potential neural sources (see (**Figure 1D, hypothesis 2**). In one case, we propose a common neural source (e.g. horizontal hMT+ cluster during horizontal perceptual state) feedbacking to both retinotopic clusters in V1 by an excitatory feedback to the matching cluster and an inhibitory feedback to the not-matching one. As observed in **Figure 4C-D**, both feedback signals in V1 are characterized by similar temporal profiles. We highlight that despite efforts from (Moon et al., 2021; Poplawsky et al., 2015) the contribution of feedback signal modulating inhibitory or excitatory neurons to BOLD fMRI at the laminar level is still unknown. In the other case, we propose that each hMT+ cluster feedbacks only the excitatory neurons belonging to the homologous retinotopic cluster in V1. Despite stronger activity of the matching hMT+ cluster, the not-matching hMT+ cluster shows only a modest reduction in activity and could, thus, be the source of feedback to the corresponding V1 clusters (see arrow from non-perceived (blue) vertical motion cluster to blue V1 (**Figure 4**). More generally, the temporal dynamics of columnar clusters in hMT+ in both the physical and ambiguous condition (**Figure 4**) resemble competitive behavior at the neural level between horizontally and vertically preferred neurons that compete to resolve the ambiguity of the stimulus: both clusters are initially activated and, during this temporal phase, are responsible for local feedback to the homologous retinotopic cluster in V1. Even if, as reported Evers et al. (2023) the neural competition is expected at much faster temporal scale (milliseconds), our fMRI results might still be valid due to the hemodynamic coupling that biologically add a 4-6 s delays to the neural response. We assume that hMT+ is the main driver in resolving the motion ambiguity and sending feedback signals to V1. However, due to the limited field of view, we can’t exclude the involvement of higher order areas (e.g. prefrontal cortex). Similarly to Liu et al. (2019), future studies with larger coverage (but likely lower resolution), could help understanding how bistable perception is resolved in the entire human brain and which areas and neural dynamics are required for that.

### 3.2 Limitations and future directions

#### Experimental paradigm

The temporal structure of the ambiguous motion condition presents significant challenges for this study. During the experiment, participants switch their perception at different times, creating an additional variability compared to the physical motion condition. A possible solution to control for it would be to match the temporal dynamic between the two conditions, by replaying the perceptual alternation of the ambiguous condition (subject’s switching times) as physical motion condition. In this way, each trial duration will have its matching control trial. However, short switches will still be discarded, reducing the statistical power in both conditions. The absence of a baseline condition between the horizontal and vertical motion conditions further complicates the interpretation of the event-related response during the ambiguous condition. Based on our results, a complete return to baseline between perceptual states would provide clearer insights on the functional response of V1 retinotopic clusters during each state separately. An alternative design for the ambiguous condition could initiate a rest condition after each perceptual state. Additionally, the absence of eye-tracking is a limitation. Without eye-tracking data, we cannot ensure that participants maintained steady gaze during the ambiguous condition, potentially confounding the retinotopic mapping in V1. However, based on the report by (Schneider et al., 2019), by choosing participants with steady fixation evaluated during behavioral assessment, this effect can be mitigated.

#### BOLD laminar profiles and draining vein effect

Even though in the last decade (2013-2024), research questions addressed with layer-fMRI have increased nearly eightfold https://layerfmri.com/, common consensus has not yet been reached regarding which acquisition and analysis methods to use. The choice of the sequence (e.g. GE, SE, VASO) together with the contrast mechanism (e.g. BOLD, CBV, CBF, CMRO2) is strictly tied to the design of the experimental paradigm and results as a compromise between signal sensitivity, specificity, temporal resolution, spatial resolution and brain coverage (Bandettini et al., 2021; Goense et al., 2016; L. Huber et al., 2019; Koopmans and Yacoub, 2019; Norris and Polimeni, 2019). Our choice of BOLD fMRI data at 0.8 iso mm fMRI using GE-EPI with 2D readout (Moeller et al., 2010) was based on a good trade-off between high sensitivity (especially for the ambiguous condition), temporal resolution (TR=2 s to accommodate the sampling of the dynamics of the switches in motion perception during the ambiguous motion condition, which mostly ranges from every 3 s to 10 s, see Supplementary Figure 1) and extensive coverage (58 slices to image both V1 and hMT+), at the cost of a reduced spatial specificity. This is due to the “draining vein effect”, a well-known phenomenon for which the venous blood carried upwards through ascending cortical veins (macrovascular weight) introduces a spurious signal which increases from deep to superficial layers that can mislead the detection of the correct neuronal sites, more coupled with capillaries and venules (microvascular weight). To mitigate this effect, we included in our experimental design both the physical and the ambiguous conditions: under the assumption that the macrovascular weight linearly sum up to the microvascular one, the subtraction operation between multiple conditions can isolate the laminar difference of interest (Aitken et al., 2020; Fracasso et al., 2018; Heynckes et al., 2023; van Mourik et al., 2023). As a consequence, differential laminar profiles can be interpreted under this simplification, as reported in 3. Alternative approaches can be used to correct for the draining vein bias https://layerfmri//unwanted vein bias. Among these, deconvolution models are considered the most physiologically plausible (Havlicek and Uludăg, 2020; Heinzle et al., 2016; Markuerkiaga et al., 2021; Marquardt et al., 2018), as they model the BOLD signal in each layer as a mixture of neural activity from multiple layers. However, higher complexity comes with the price of additional assumptions about the underlying vascular architecture, neurovascular coupling and interlaminar leakage. This can lead to overly complicated interpretations of laminar results, particularly when multiple cortical brain areas are involved, as is the case in our study. Moreover, the parameters of the model set to mimic a conventional draining veins effect observed mainly in feedforward dominated condition, might not be optimal for our feedback condition.

Laminar results on feedback conditions in V1 seems to be strongly dependent on the experimental paradigm: superficial layers of V1 have been found to reflect feedback in contextual filling-in research using the occlusion paradigm (Muckli et al., 2015) and visual perception (Bergmann et al., 2024). Contrary, static illusion e.g. Kanizsa (Kok et al., 2016; Marquardt et al., 2020) or working memory tasks (Lawrence et al., 2018) showed a modulation in deep layers. Although the variety of the laminar results in V1 could indicate a discordance in the field, it has to be acknowledged that different feedback pathways from higher-areas to V1 do co-exist (Callaway, 2004). Using laminar-fMRI, Bergmann et al. (2024) demonstrate that different feedback pathways in V1 are recruited according to the cognitive task: while their mental imagery tasks elicit modulation in deep layers, illusory visual perception tasks target superficial layers.

### 3.3 Challenges of layer-fMRI

Although layer-fMRI is currently the most promising technique for non-invasively investigating the concurrent mesoscopic organization of multiple cortical areas (such as V1 and hMT+ in our study), several limitations ought to be acknowledged.

One significant challenge is the unknown translation of the oscillatory behavior of the neurons to the coupled fMRI amplitude at the laminar scale, particularly since feedback and feedforward processes are associated with different frequency bands (Bastos et al., 2015; Spyropoulos et al., 2018). Layer-fMRI studies operate under the assumption that hemodynamic coupling (energy requirement) remains the same despite these differences in frequency coupling.

Cognitive processes as bistable perception have been modeled at neuronal scale and validated by electrophysiology data as continuous exchange of information occurring at much higher temporal scales (Brascamp et al., 2015; Cao et al., 2021; Evers et al., 2023; Moreno-Bote et al., 2010; Shpiro et al., 2007). While using the hemodynamic correlate of neural activity provides a wider coverage compared to electrophysiology experiments, the resulting fMRI activity may only represent the tip of the iceberg compared to the underlying neural processes.

Additionally, it is generally assumed that the BOLD fMRI response in the cortex primarily reflects changes in excitatory neural activity, as excitatory neurons constitute 80–90% of all cortical neurons (Meyer et al., 2011). However, the ratio between excitatory and inhibitory neurons varies across cortical depth and areas (Markram et al., 2004; Tremblay et al., 2016), and its relationship with hemodynamic coupling remains unknown, potentially biasing the interpretation of layer-fMRI results (Moon et al., 2021). For instance, our experimental design using the motion quartet allows us to separately study the functional role of each cluster during both its expected preferential and non-preferential motion conditions. The positive signal change observed during both conditions in V1 retinotopic clusters (**Figure 4C**) is difficult to interpret. It could indicate that a feedback signal is received and processed by both cluster types. The cluster corresponding to the perceived motion might be enhanced by excitatory feedback, while the opposite cluster might be suppressed by inhibitory feedback. However, these two fMRI signals, which appear indistinguishable at the current cutting-edge spatio-temporal scale of layer-fMRI, might conceal more complex dynamics.

Generative neural models can complement empirical results with mechanistic interpretation that are generated at neural level (Friston et al., 2003; Havlicek et al., 2017; Uludag and Havlicek, 2021). However, even these methods are limited by the fact that the same neuronal excitation-inhibition coupling model is used across different layers and across cortical brain areas. More realistic models of the underlying angioarchitecture are also expected to complement our understanding of the neurovascular coupling. Current efforts in pushing the in-vivo anatomical MRI resolution up to 0.35 iso mm (Gulban et al., 2022) within a reasonable scanning time, multimodal studies on combining post-mortem microscopy and quantitative MRI in humans (Alkemade et al., 2022), in advancing in vivo histology together with biophysical models (Dinse et al., 2015; Weiskopf et al., 2021) are crucial for improving our understanding of the mechanistic principle of layer-fMRI and close the gap with the electrophysiology data.

### 3.4 Conclusion

In conclusion, this study enhances our understanding of the neural mechanisms underlying bistable perception in humans, particularly the interaction between early and high-level visual areas in the brain. By utilizing ultra-high-field layer-fMRI, we demonstrated that hMT+ plays a crucial role in resolving perceptual ambiguity through feedback mechanisms that modulate activity in V1. A global feedback mechanism from hMT+ to both retinotopic clusters of V1 is reflected in the increased similarity in temporal dynamics between the two areas of interest. This research not only clarifies the role of feedback in visual perception but also provides a crucial foundation for further exploration of the neural dynamics at the mesoscopic scale that support conscious experience.

## 4 Methods

### 4.1 Experimental design

#### 4.1.1 Participants

Nine healthy participants (7 females and 2 males) with normal or corrected-to-normal vision were recruited for the study. Participants received a monetary reward. All participants had been in an MRI scanner at least once before and been trained and experienced at maintaining fixation for long periods of time. Informed consent was obtained from each participant before conducting the experiment. The study was approved by the ethics review committee of the Faculty of Psychology and Neuroscience (ERCPN) of Maastricht University and experimental procedures followed the principles expressed in the Declaration of Helsinki.

#### 4.1.2 Pre-scan behavioral session

The perception of motion elicited during ambiguous stimulation can vary substantially across participants: individuals might switch between horizontal and vertical motion perception very rapidly (every 1-2 seconds), but they may also perceive one type of motion (e.g. vertical) more frequently than the other (Chaudhuri and Glaser, 1991). In order to maximize the efficiency of sampling perceptual responses during fMRI acquisition, we selected our participants according to the results of a behavioral pre-scan session. We invited potential participants to collect three runs of ambiguous motion data over 1h. We chose participants who experienced a stable switch between horizontal and vertical motion (or vice versa) occurring at least once every 5 seconds throughout the run.

#### 4.1.3 Stimulus description

Each participant underwent two (f)MRI sessions on two separate days. Each session lasted for 2h. During the first session, we collected 1 run to functionally locate hMT+, 5-7 runs of the motion quartet stimulus by alternating physical and ambiguous versions, and an MP2RAGE scan to obtain a high-resolution structural image. During the second session we collected 5-7 runs of motion quartet stimulus with the same alternation scheme (for a total of 6 runs for the physical and 6 runs for the ambiguous quartet across two sessions) and two runs of population receptive field mapping.

##### hMT+ functional localizer

A standard block design paradigm presenting moving dots in alternation with static dots in a circular aperture (Tootell et al., 1995) (run duration: 9 min 10 s) was used to cover the whole human motion complex (hMT+), without separating MST from MT), as previously described (Huk et al., 2002; Kolster et al., 2010; Pizzuti et al., 2023; Zimmermann et al., 2011). Briefly, dots traveled inwards and outwards from the center of the aperture for 10 s (speed = 8 degree of visual angle per second, dot size = 0.2 degree of visual angle, number of dots = 200, black dots on gray background), were followed by a stationary dots display presented for the same amount of time. A total of 14 repetitions of task-rest blocks were collected. The duration of this run was 4 min 50 s (270 volumes, TR = 1 s).

##### The motion quartet

We presented the motion quartet stimulus inducing a similar illusory horizontal and vertical motion as the presented physical horizontal and vertical motion stimuli (**Figure 1A**): the physical motion and the ambiguous quartet were designed as described in Schneider et al. (2019). The quartet is composed of four squares (1°x1° visual angle) so-called ‘inducers’ whose horizontal distance was set to 3° and vertical distance to 3.7° (Schneider et al., 2019). In the physical quartet, squares moved along the horizontal or vertical paths for 10 s per condition. After four alternations of horizontal and vertical motion (80 s), a flicker condition where all inducers synchronously blink (16 s) was shown to serve as baseline. This local flicker stimulus did not induce a motion percept. This scheme was repeated six times. Each run started and ended with a simple fixation condition (lasted for 20 s). In the ambiguous quartet, while keeping the same condition scheme of the physical quartet, we replaced the horizontal and vertical motion with a constant ambiguous quartet stimulus (80 s) that induced perception of horizontal or vertical apparent motion through blinking squares at diagonally opposite corners (80 s). A pair of squares was presented for 150 ms (9 frames) followed by an inter-stimulus interval of 67 ms (4 frames). Such a presentation frequency of 2.3 Hz was shown to induce strong perception of apparent motion (Finlay and von Grünau, 1987; Schneider et al., 2019). Participants indicated percepts via a MR-compatible button box (whether button ”1” or button ”2” was used to indicate horizontal or vertical motion). The duration of each motion quartet run was 10 min and 30 s (309 volumes, TR = 2 s)

##### Retinotopic mapping stimulus

For population receptive field mapping we used a bar aperture (0.9° wide) revealing a flickering chromatic checkerboard pattern (4 Hz) that was presented in 4 orientations (−45°, +45°, 0°, 90°). For each orientation, the bar covered the entire screen (−5° to 5° visual angle) in 12 discrete steps (each lasting 1 s). Within each orientation, the sequence of steps (and hence the locations) was randomized and each orientation was presented 6 times. Our stimulus presentation combined the chromatic contrast approach from Swisher et al. (2007) and the bar based approach from Dumoulin and Wandell (2008). The duration of the population receptive field mapping run was 5 min and 13 s (309 volumes, TR = 1 s). Similar to the main experiment, subjects were instructed to perform a central fixation task during the retinotopic mapping experiment and respond through the button box every time the fixation dot changes its color (attention task).

##### Stimulus presentation

The stimulation scripts were presented using the open source application PsychoPy3 (v2020.2.4). Scripts are available on https://github.com/27-apizzuti/meso MotionQuartet/ stimulus scripts. A frosted screen (distance from eye to screen: 99 cm; image width: 28 cm; image height: 17.5 cm) at the rear of the magnet was used to project the visual stimuli (using Panasonic projector 28 PT-EZ570; Newark, NJ, USA; resolution 1920 x 1200; nominal refresh rate: 60 Hz) that participants could watch through a tilted mirror attached to the head coil. We used 50% gray background (at 435 cd*/*m2 luminance) with white dots (at 1310 cd*/*m2) for the motion stimulation (black color is measured at 2.20 cd*/*m2).

### 4.2 MRI data acquisition

Data acquisition was performed on a whole-body “Plus” MAGNETOM 7T (Siemens Healthineers, Erlangen, Germany) at Scannexus B.V. (Maastricht, The Netherlands) using a 32-channel RX head-coil (Nova Medical, Wilmington, MA, USA). The shimming procedure included the vendor-provided routines to maximize the field homogeneity within the imaging slab. We found that the shimming was quite effective (spatially varying off-resonance frequencies occur on a very large spatial scale). The hMT+ localizer and retinotopic mapping experiment was conducted using a 2D GE EPI sequence with BOLD contrast (Moeller et al., 2010) as previously done in Pizzuti et al. (2023) with some adaptations in imaging parameters, (echo time (TE) = 23 ms, nominal flip angle (FA) =54°, echo repetition time (TR) = 1000 ms, multi band factor (MB) = 3, 48 slices) with an almost whole brain field of view and a (100 × 100) matrix at 1.8 mm isotropic nominal resolution. For the motion quartet experiment we used 2D GE EPI sequence with BOLD contrast (Moeller et al., 2010) with 0.8 isotropic mm resolution with partial coverage. The in-plane field of view was 140×137 mm (176×172 matrix) for a total of 58 acquired slices. The imaging parameters were: TE = 24.6 ms,TR = 2000 ms, flip angle FA = 69°, in plane partial Fourier factor 6/8, GRAPPA=3, MB=2. The placement of the small functional slab during the first session was guided by an online analysis of the hMT+ localizer data (general linear modeling by Siemens), to ensure a bilateral coverage of early visual areas and hMT+ for every participant. Before the acquisition of the run, we collected 5 volumes for distortion correction with the settings specified above but opposite phase encoding (posterior-anterior). An auto-align (AA-scout) sequence was used to assure that the determined field of view was placed in the same position across sessions. The anatomical images were acquired with an MP2RAGE (magnetization prepared 2 rapid gradient echoes) (Marques et al., 2010) at 0.7 mm isotropic resolution (TR/TE = 6000 ms/2.39 ms, TI = 800 ms/2750 ms, FA = 4°/5°, GRAPPA = 3). MP2RAGE sequence parameters are optimized to overcome the large spatial inhomogeneity in the transmit B1 field by generating and combining in a novel fashion 2 different images at 2 different inversion times (TI1, TI2) to create T1-weighted MP2RAGE uniform (UNI) images (Marques et al., 2010).

### 4.3 Anatomical data analysis

#### 4.3.1 Preprocessing and surface reconstruction

T1-weighted UNI images with high contrast-to-noise ratio from MP2RAGE were used to reconstruct cortical surfaces and derive layers in the cortical ribbons of interest (V1, hMT+). T1-w UNI images were skull-stripped using a brain mask obtained by inputting the MP2RAGE INV2 (TI2) images to FSL BET (v.6.0.5) (Smith et al., 2004) and corrected for intensity inhomogeneities using N4BiasFieldCorrection (Tustison et al., 2010). We upsampled our anatomical images at 0.35 isotropic mm resolution and we reconstructed cortical surfaces by following the ‘advanced segmentation pipeline’ in BrainVoyager (Goebel, 2012) with few adaptations. As a preliminary step, we created a subcortical mask in Brainvoyager at the recommended resolution of 0.5 isotropic mm resolution and in Talairach space. The usage of a standard space with known reference points is beneficial to compute the mask. Using the ‘c3d reslice’ program from ITK-SNAP (Yushkevich et al., 2006) we resampled the mask in the native space at 0.35 isotropic mm resolution and applied it to T1-w UNI images before undergoing the advanced segmentation pipeline. Following the pipeline, we generated whole-brain white and gray matter tissue segmentation that was, afterwards, manually corrected in ITK-SNAP with a particular focus around the calcarine and the middle temporal sulcus. Connected cluster detection algorithm, one/two iterations of tissue regularization with morphological operations, manual edits to remove remaining bridges, were applied before reconstructing white matter surfaces for both hemispheres in BrainVoyager. Each surface was smoothed, downsampled to 160k vertices and mapped to a standard sphere using curvature-driven cortex-based alignment (CBA) in BrainVoyager. Our regions of interest (V1 and hMT+) were manually drawn in the surface space (see ROI definition paragraph) and projected back to the volume space (sampling the gray matter from white matter surface 0 to +2 mm for V1 and from 0 to +3 mm for hMT+).

#### 4.3.2 Volumetric cortical depth sampling

Cortical depth sampling was performed in the native volumetric space (T1-w UNI) at 0.35 mm x 0.35 mm x 0.35 mm for each subject by using LayNii software (v2.2.1) (Gulban et al., 2022; L. (R. Huber et al., 2021). We restricted the initial whole-brain white and gray matter segmentations from the previous step, to a focal mask that encapsulates our ROIs (this mask was created by dilating 3 times the volume of our combined ROIs). Tissue labels were carefully quality controlled and manually edited in ITK-SNAP when necessary (by A.P.), and later revised independently by another expert (O.F.G.). Finally, we ran the LN2 RIM POLISH program from LayNii (based on morphological operations) to polish the result of our manual segmentation. We propagated the definition of V1 and hMT+ ROIs to the final gray matter segmentation using the LN2 VORONOI program from LayNii. Note that to restrict the propagation to the radial axis of the gray matter, we added a ‘capsule’ ROI for V1 and hMT+ also defined in the surface space. We used the LN2 LAYERS program from LayNii to compute equi-volume cortical depths (D coordinate) (Bok, 1959; Waehnert et al., 2014), cortical thickness and curvature for each gray matter voxel. The equivolume layers are used to show our laminar results (see Supplementary Figure 4).

#### 4.3.3 Cortical volumetric parametrization

We obtained a full volumetric parametrization around V1 and hMT+ restricted to a sub-volume that fully encapsulate significant functional response obtained from the physical motion condition (see horizontal and vertical clusters described in ‘ROI definition’ paragraph) by using the LN2 MULTILATERATE program from LayNii (Gulban et al., 2022). This program computes a pair of orthogonal coordinates (U,V) for a disk of interest with a predetermied radius that geodesically grows from an initial ‘control point’ set by the user (usually this control point is placed in the center of an activated region). The radius also varies across ROIs according to its curvature and to the spatial extent of the functional activity. U, V coordinates together with D coordinate resulting from the cortical depth parametrization (LN2 LAYERS-equivol) fully parameterized the volume of interest.

### 4.4 Functional data analysis

#### 4.4.1 Preprocessing and registration

All fMRI data (hMT+ localizer, motion quartet and pRF experiment) underwent the same preprocessing steps: slice time correction (BrainVoyager), motion correction (BrainVoyager), correction for geometric distortion (fsl-topup, Smith et al., 2004, high-pass filter (BrainVoyager) (3 cycles for hMT+ localizer and pRF experiment, 5 cycles for motion quartet as previously done by Schneider et al. (2019). The target volume for the motion correction was chosen for each experiment type as the first volume of the first run within the same scanning session. One subject was excluded from further analysis since motion parameters exceeded 1.6 mm (double of voxel resolution) for 4 runs of motion quartet. A pair of opposite phase encoding images were acquired in each session at the beginning of the first functional run of each experiment type and used to estimate the susceptibility-induced off-resonance field and correct for geometric distortions. Boundary-based registration (BBR) as implemented in BrainVoyager was used to align preprocessed fMRI data from each experiment type and session to the anatomical space T1w UNI images. Note that for across-sessions alignment an initial grid search approach was used to improve the co-registration (BrainVoyager). Goodness of alignment was assessed with the quantitative cost function computed in BrainVoyager being below 0.3 for fMRI at 1.8 iso mm3 and 0.1 for 0.8 iso mm3 and with a qualitative inspection. We used the anatomical space at 0.7 isotropic mm resolution as target space for all functional runs. This approach with series of linear transformation was chosen over using alternatives e.g. choosing one functional run as target space in combination with non-linear transformation (for example as done by Pizzuti et al. (2023)) for its flexibility with regards to anatomical processing steps (using ‘distortion free’ images to reconstruct surfaces, depth sampling), and to multi-sessions and multi-scale fMRI data alignments.

#### 4.4.2 Region of interest definition

Bilateral hMT+ was functionally defined based on the results of a voxel-wise general linear model (GLM) computed for the localizer run. The GLM computed in BrainVoyager contained a single predictor for the stimulus condition “moving dots” convolved with a standard hemodynamic response function (see Pizzuti et al., 2023). The model was corrected for temporal auto-correlation (AR2). Statistical maps (t-map) were projected on the reconstructed white matter surface (integrating t-values along vertex normals from 0-to-3 mm into the gray matter. Estimated hMT+ thickness is 3 mm) and ROI boundaries were manually drawn encapsulating voxels whose functional response to “moving vs static dots” contrast was significant (using a threshold (q) corrected for multiple comparisons using false discovery rate; q(FDR) *<* 0.05) and contained in one distinct cluster (Schneider et al., 2019). Surfaces ROIs were projected back into the volume space following the same integration scheme (0-3 mm into the gray matter). Early visual areas (V1, V2, V3) from both the dorsal and ventral pathway were manually delineated using polar angle and eccentricity maps projected on reconstructed white matter surfaces (integrating values along vertex normals from 0-to-2 mm into the gray matter. Estimated V1 thickness is 2 mm). Volumetric maps are computed using the population receptive field mapping approach to retinotopy, where receptive fields are modeled as parametric forms (e.g., isotropic Gaussian), as described in (https://github.com/ ccnmaastricht/CNI toolbox/wiki/Population-Receptive-Field-Mapping-(Python))(Senden et al., 2014). We draw our ROIs according to the eccentricity and polar angle maps (maps were thresholded according to *R*2 *>* 0.2, indicating the goodness of fitting) following polar angle reversals and orthogonality with respect to iso-eccentricity lines (Dumoulin and Wandell, 2008; Sereno et al., 1995). One smoothing iteration was performed on surface-projected eccentricity maps. Finally, ROIs were projected back into the volume space following the same integration scheme (0-2 mm into the gray matter). Note that, we limited our analysis to V1 since the definition of V1/V2 borders was straightforward for all participants.

#### 4.4.3 Univariate statistical analysis and definition of mesoscopic functional clusters

To investigate univariate differences between ROIs (V1 and hMT+), we computed a multi-run voxelwise GLM analysis for both physical and ambiguous conditions. Time course was normalized within BrainVoyager according to *y norm* = *y/y mean ∗* 100. The GLM was corrected for temporal auto-correlation (AR2). We computed separate GLM per condition (physical and ambiguous) since we expect the two conditions to elicit different brain responses. Three predictors (horizontal motion, vertical motion and flicker) were modeled with canonical (two-gamma) hemodynamic response functions (HRF), time-locked to the presentation of the stimuli in the physical runs and to the subject’s perceptual responses in the ambiguous runs. Betas values for each predictors were computed and converted as percent signal change for both physical and the ambiguous condition. To assess voxel’s selective response to motion type, we computed a t-map for the contrast: horizontal *>* vertical. The two contrasting conditions were balanced in terms of the number of time points considered in the computation. At mesoscopic level, both V1 and hMT+ shows a characteristic functional organization in response to physical motion: V1 differentiates horizontal and vertical clusters according to its retinotopic organization, whereas hMT+ is comprised by axis-of-motion (or direction-of-motion on a more fine scale) functional columnar clusters (Pizzuti et al., 2023; Schneider et al., 2019; Zimmermann et al., 2011) where voxels are organized in patches according to their preferential directional motion response. Therefore, we defined V1 and hMT+ horizontal and vertical clusters by considering voxels whose functional response was highly selective during the physical motion condition (t-value for the contrast horizontal *>* vertical was higher than 95 percentile) within the defined hMT+ and V1 ROI from the independent localizer runs.

#### 4.4.4 Clusters characterization

We used the percent signal change measure from the betas estimates from GLM to characterize each mesoscopic cluster to both the horizontal and the vertical trials within both physical and ambiguous conditions. Although the physical condition is used to define horizontal and vertical clusters within V1 and MT+, we still include this condition in all the following analyses since it serves as a baseline to compare the functional modulation elicited by the ambiguous condition. First, we quantified, for each participant, the amplitude of the cluster’s response for both the physical and the ambiguous condition during the expected preferred condition by averaging the percent signal change across all voxels belonging to the cluster (e.g. for the horizontal cluster, we averaged the percent signal change relative to the horizontal condition). We then obtain a group statistic by computing the mean and the standard error of the mean estimates across participants (N=8). Scipy routine in python was used for this purpose (Virtanen et al., 2020). Results are summarized in (**Figure 2**). Second, we computed a voxel-wise measure of specificity as introduced in Pizzuti et al. (2023), Eq. 2 using percent signal change estimates, to complement the “winner takes all” approach used to associate each voxel to the horizontal or vertical clusters according to the maximum response elicited during the physical motion condition. This quantification is applied to the ambiguous motion condition. Specificity measure was averaged across voxels within the same cluster, and as previously explained, converted in group statistics (mean +/-standard error of the mean) as reported in (**Figure 4A-B**).

#### 4.4.5 Laminar analysis

In order to investigate depth-dependent features we extended V1 and hMT+ physical clusters to cover the entire cortical thickness by using the LN2 MULTILATERATE -max program from LayNii as previously applied and described in Dresbach et al., 2024; Pizzuti et al., 2023. This program makes use of a moving cylinder (radius=0.39, height=local thickness) within our volume of interest (fully parameterized by U, V, D coordinates) and propagates the cluster mask to cover the local cortical thickness even when only one voxel is activated along the depth. Laminar profiles are computed for each subject, ROI (V1, hMT+), cluster type (horizontal and vertical) and condition (physical, ambiguous) using voxel-wise betas estimates from GLM results, expressed in percent signal change. Laminar profiles are averaged within cluster type and across participants and reported as group statistics in (**Figure 3**). Due to the choice of the BOLD contrast mechanism for our fMRI data (see Discussion), a draining vein effect is known and expected to affect our laminar profile. When only one condition is considered at the time, this unwanted effect is manifested with an increasing trend from deep to superficial layers due to large ascending veins running through the cortical depth and pial veins. With the assumption of linearity between the microvascular and the macrovascular weight of the laminar BOLD fMRI signal, the computation of differential laminar profiles (physical-ambiguous) serves as mitigation of this draining vein effect. The resulting laminar signal is supposed to highlight the differences between the two experimental conditions, given the same macrovascular signal. This strategy has been widely employed within the layer-fmri literature (Aitken et al., 2020; Fracasso et al., 2018; Heynckes et al., 2023; van Mourik et al., 2023).

#### 4.4.6 Event related averages

We investigated the temporal response pattern of our selected voxels within V1 and hMT+ horizontal and vertical clusters for both physical and ambiguous conditions. For each run, we obtain a cluster time series by pulling and averaging time courses of voxels belonging to the same cluster. Trials were separated according to the condition type: horizontal and vertical. Then, each trial (t) was converted in percent signal change as follows *t* = (*t − t*(0)*/t*(0)) *∗* 100. Trials of the same types were temporally aligned to the onset time (reported as zero time). While the duration of both horizontal and vertical condition was fixed to 10 s during the physical motion condition, in the ambiguous condition epochs duration subjectively vary within a wide range (3-20 s, see Supplementary Figure 1). We aimed to match the time duration of the ambiguous trials to the physical ones, by excluding perceptual trials whose whole duration was shorter than 10 s (showing 5 TR in total). Then, we averaged the normalized time course of the remaining selected trials. This trial selection strategy did not assure the same duration for the ambiguous trials. In order to consider the same number of trials per time point for a balanced average, we were limited to display and interpret the resulting event-related averages up to 5 TR. This means that trials with duration longer than 10 s were cut to 5s. Finally, since we did not expect differences between the two motion cluster types, we summarized our results by averaging event-related responses from both clusters in their ’preferred’ condition (horizontal trials for the horizontal cluster, vertical trials for vertical cluster) and ‘not preferred’ condition (horizontal trials for the vertical cluster, vertical trials for horizontal cluster). Averaging at multiple scales helped to boost the signal especially for the ambiguous condition where multiple trials had to be discarded to obtain a uniform duration. Group statistics reported as mean and standard error the mean within our sample (N=8) are shown in **Figure 4C-D**.

## 5 Data and Software availability statement

Analysis code is available on GitHub: https://github.com/27-apizzuti/meso MotionQuartet. Raw data are shared in Zenodo: https://doi.org/10.5281/zenodo.13737438, https://doi.org/10.5281/zenodo.13738292, https://doi.org/10.5281/zenodo.13739150 Please note that, due to the upload size limit, we have divided our dataset into three parts.

## Acknowledgements

This project was funded by the EU-project H2020-860563 euSNN and the European Union’s Horizon 2020 Framework Programme for Research and Innovation under the Specific Grant Agreement No. 945539 (Human Brain Project SGA3). Data was acquired at Scannexus (Maastricht, the Netherlands). Laurentius Huber was funded from the NWO VENI project 016.Veni.198.032. OFG is funded by Brain Innovation. We thank Federico de Martino for helping with setting the scanning sequence. We thank Alexander Kroner for the population receptive field stimulus. We thank Kris Evers for helpful discussions on the laminar dynamics. We thank Logan Dowdle and Jesse Breedlove for relevant discussions on the limitation of the current experimental paradigm. Finally, we thank “Maastricht layer-seminar” members Sebastian Dresbach, Lonike Faes, Miriam Heynckes, Kenshu Koiso, and Yawen Wang for discussions throughout the project.

## 7 Author Contributions

According to the CRediT system (https://casrai.org/credit/)

**Conceptualization:** A.P., R.G.

**Methodology:** A.P., O.F.G., R.H., J.P., R.G.

**Software:** A.P., O.F.G.

**Validation:** A.P., O.F.G., R.G.

**Formal Analysis:** A.P.

**Investigation:** A.P., O.F.G.

**Resources:** A.P., R.H., O.F.G., J.P., R.G.

**Data curation:** A.P.

**Writing – original draft:** A.P.

**Writing – review & editing:** A.P., R.H., O.F.G., J.P., R.G.

**Visualization:** A.P., O.F.G., R.G.

**Supervision:** O.F.G., J.P., R.G.

**Project administration:** R.G.

**Funding acquisition:** R.G.

## 8 Supplementary Material

**Supplementary Figure 1:**
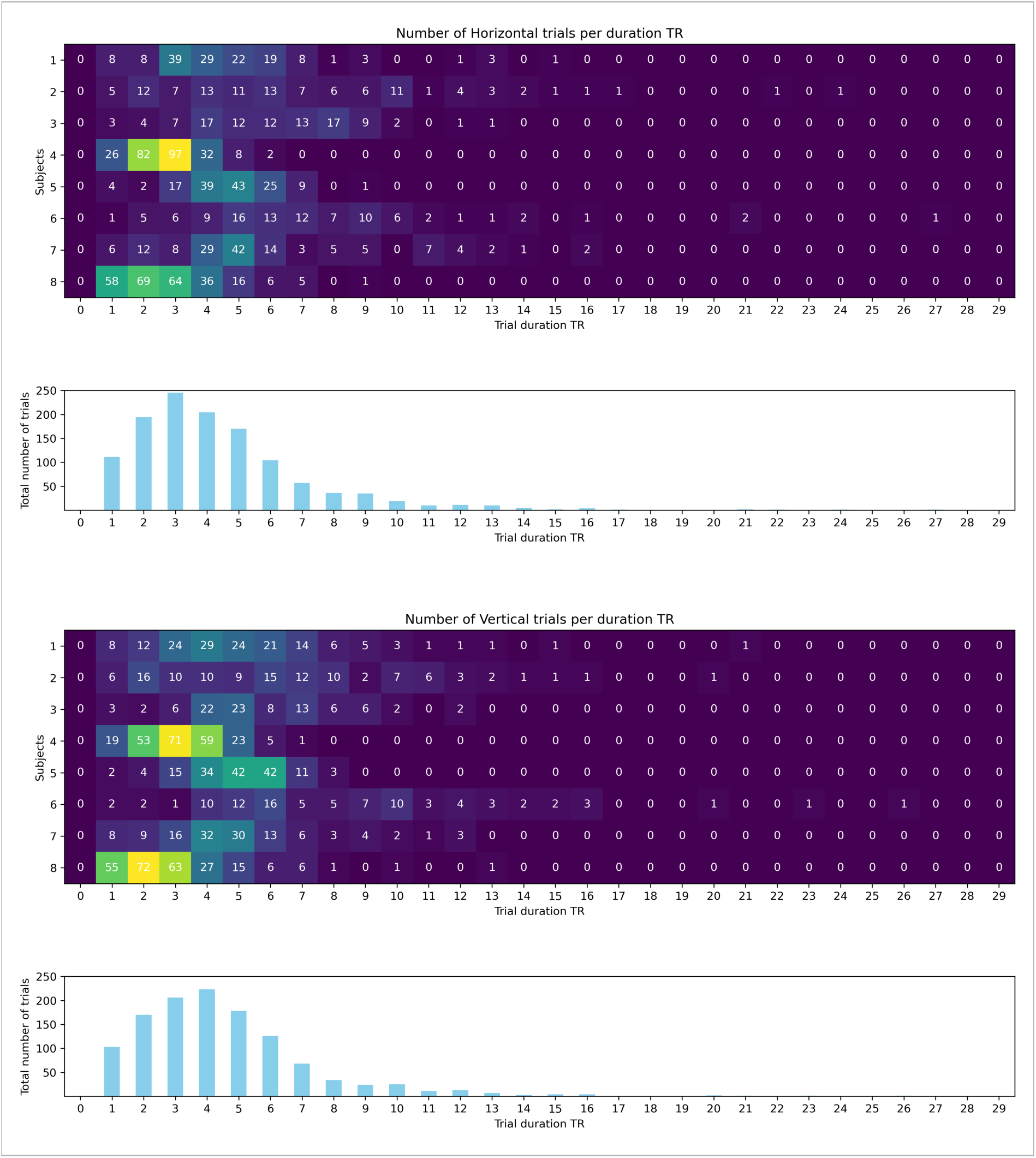
Distribution of trial durations (horizontal and vertical) during the ambiguous motion condition for all the subjects.

**Supplementary Figure 2:**
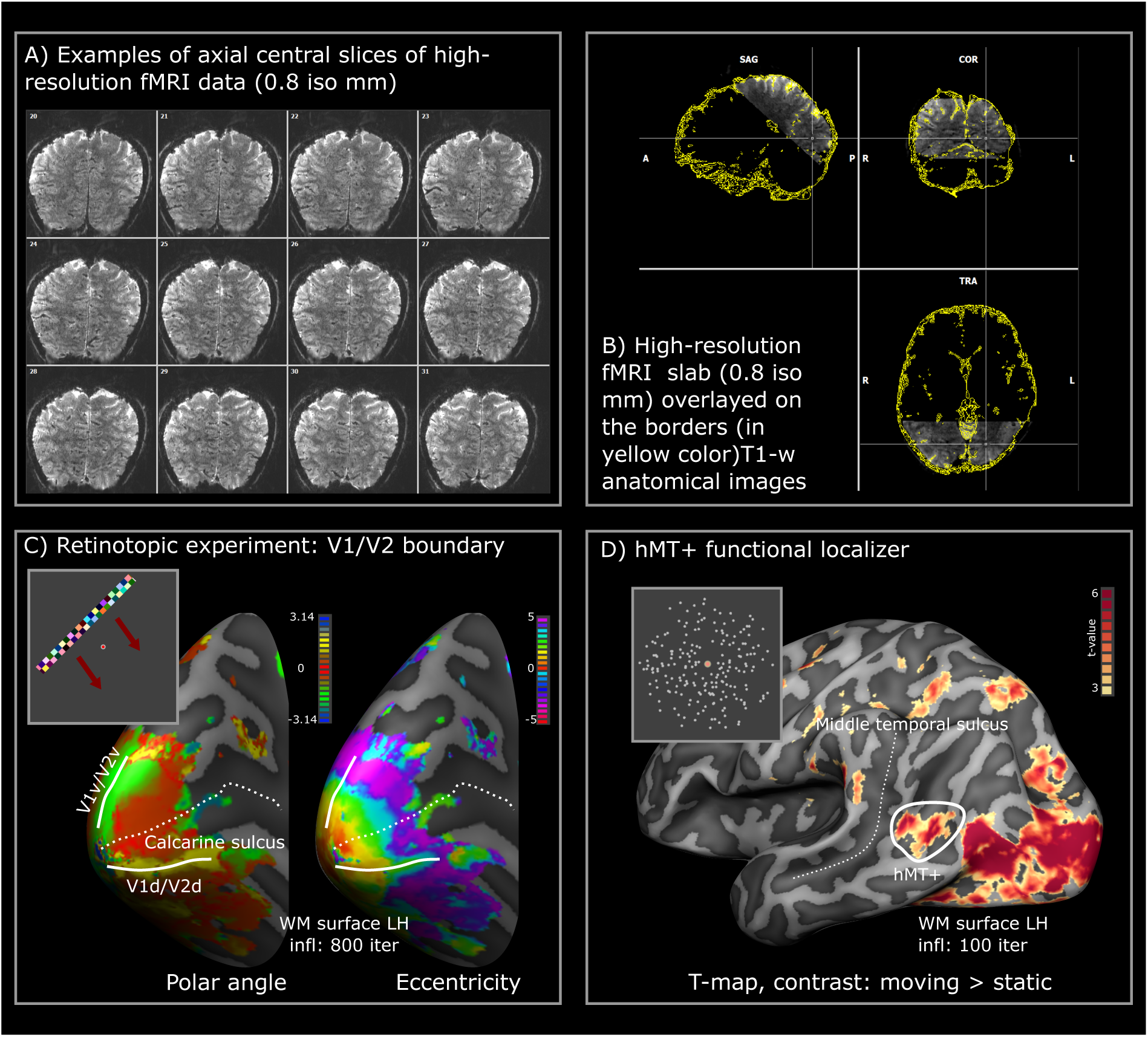
Data quality and region of interest definition (sub-04). A) Examples of fMRI slices at 0.8 iso mm. B) High-resolution fMRI coverage. C) Example of polar angle and eccentricity maps obtained during the retinotopic experiment and manually drawn V1/V2 borders (white solid lines reported on the surface). D) Statistical t-map from hMT+ functional localizer run. White circle on the surface indicate hMT+ region of interest.

**Supplementary Figure 3:**
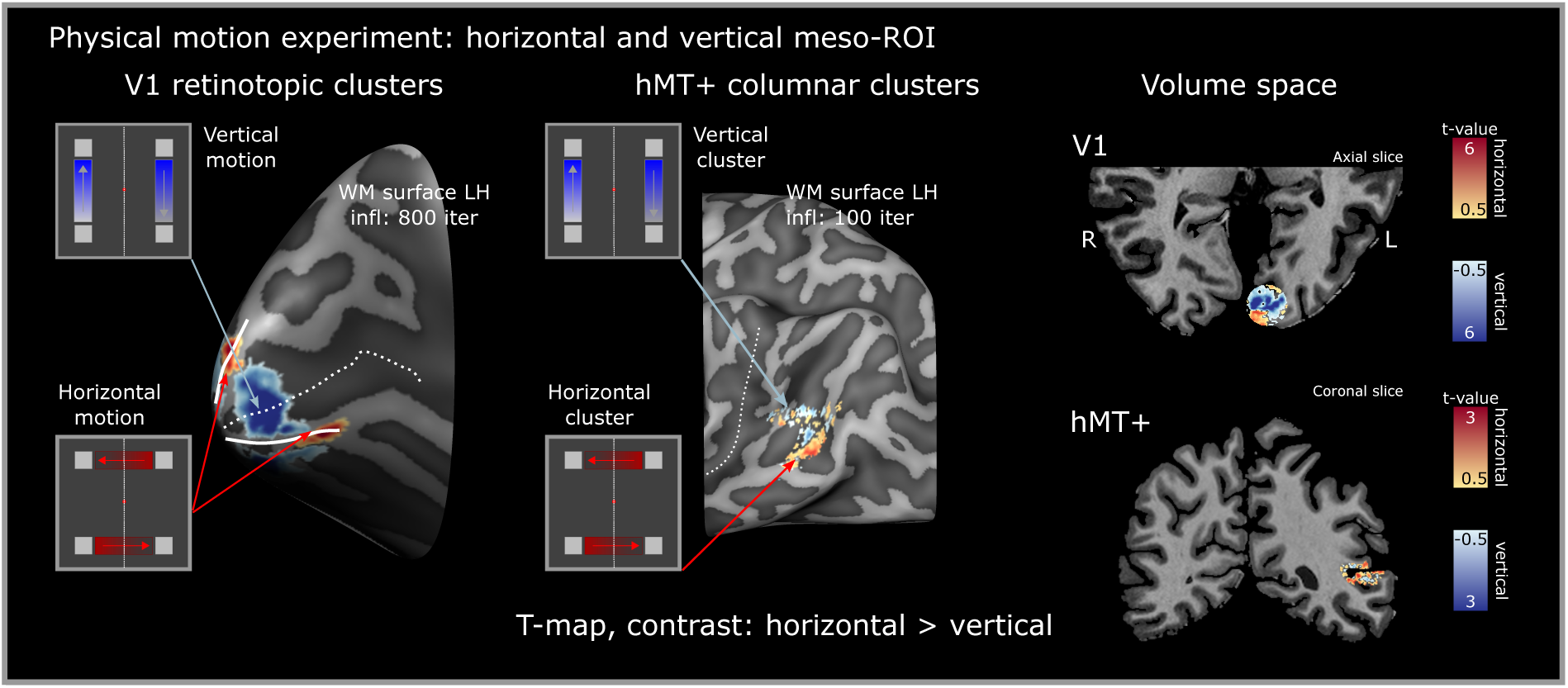
Mesoscopic cluster definition (sub-04). Statistical t-maps (contrast: horizontal *>* vertical) from physical motion runs are used to define horizontal and vertical clusters (retinotopic clusters in V1, motion-specific clusters for hMT+). Maps are shown both in the surface and volume space on one exemplary slice.

**Supplementary Figure 4:**
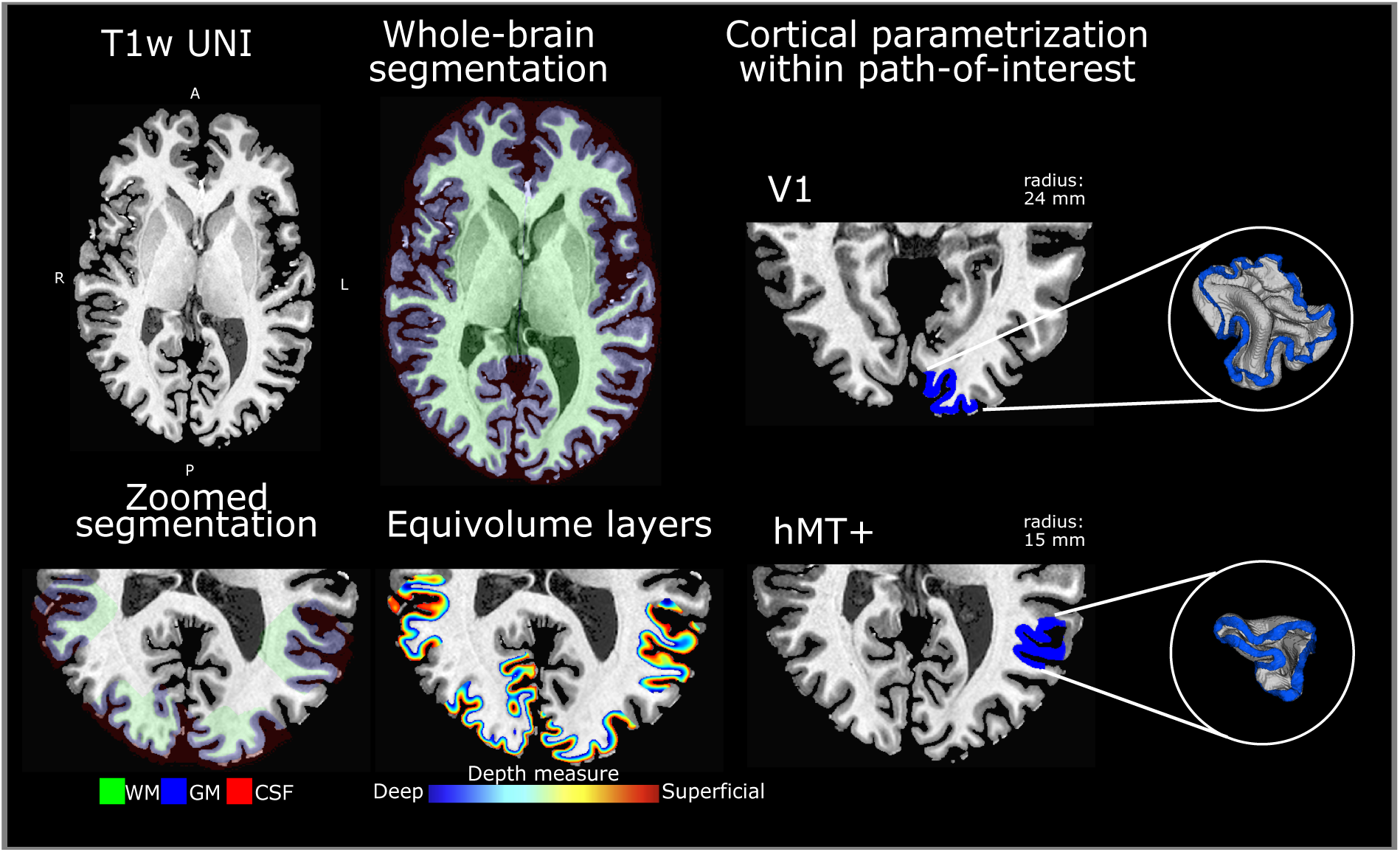
Segmentation steps (sub-04). T1w UNI image from MP2RAGE is inputted in BrainVoyager advanced segmentation pipeline to obtain an initial whole brain segmentation. After manual editing, white matter segmentation was used to reconstruct the white matter surface used for drawing the regions of interest (V1/V2 borders, hMT+). Once delineated the ROIs, we further polished tissue segmentation around them (‘Zoomed segmentation’ step). Normalized equivolume depth-coordinate D and cortical layers are computed using LN2 LAYERS -equivol in LayNii. Finally, the LN2 MULTILATERATE program in LayNii is used to generate U,V cortical coordinates that together with D coordinate provide a whole 3D parametrization (‘Cortical parametrization within patch of interest’ step).

